# Encapsulation of AAVs into protein vault nanoparticles as a novel solution to gene therapy’s neutralizing antibody problem

**DOI:** 10.1101/2023.11.29.569229

**Authors:** Logan Thrasher Collins, Wandy Beatty, Buhle Moyo, Michele Alves-Bezerra, Ayrea Hurley, Qing Lou, Z. Hong Zhou, William Lagor, Gang Bao, Selvarangan Ponnazhagan, Randall McNally, Leonard H. Rome, David T. Curiel

**Affiliations:** Washington University in St. Louis Department of Biomedical Engineering; Washington University in St. Louis Department of Radiation Oncology; Washington University in St. Louis Department of Molecular Microbiology; Rice University Department of Bioengineering; Baylor College of Medicine Department of Molecular Physiology and Biophysics; California NanoSystems Institute; University of California Los Angeles Department of Microbiology, Immunology, and Molecular Genetics; University of California Los Angeles Department of Materials Science and Engineering; Baylor College of Medicine Department of Integrative Physiology; University of Alabama at Birmingham Department of Pathology; Vault Pharma Inc.; University of California Los Angeles Department of Biological Chemistry

## Abstract

Although adeno-associated virus (AAV) has enjoyed enormous success as a delivery modality for gene therapy, it continues to suffer from the high prevalence of preexisting neutralizing antibodies in human populations, limiting who can receive potentially life-saving treatments. In this regard, AAV therapies generally also must be administered as a single dose since neutralizing antibodies develop in patients who receive the virus. Strategies for circumventing these issues remain limited. As a novel solution, we employed SpyTag-SpyCatcher molecular glue technology to facilitate packaging of AAVs inside of recombinant protein vault nanoparticles. Vaults are endogenous particles produced by mammalian cells. We therefore hypothesized that they may shield packaged molecules from neutralizing antibodies. Vaults have previously been utilized to deliver drugs and proteins into cells, but our study represents the first time anyone has packaged an entire virus inside of a vault. We showed that our vaultAAV (VAAV) delivery vehicle transduces cells in the presence of anti-AAV neutralizing serum. VAAV is positioned as a new gene therapy delivery platform with potential to overcome the neutralizing antibody problem and perhaps even allow administration of multiple doses, expanding the scope of AAV treatments.

## Introduction

Adeno-associated virus (AAV) is one of the most successful vehicles for therapeutic gene delivery. Several AAV gene therapies are available on the U.S. market, including Luxturna, Zolgensma, Hemgenix, Elevidys, and Roctavian.^1,2^ Dozens of AAV clinical trials are underway,^3^ and commercial players are investigating novel ways of engineering AAVs for greater efficacy.^1,3,4^ Despite their momentum and strong clinical profile, progress in the AAV gene therapy field has slowed in recent years due to adverse immunological responses against the vector.^3,5,6^ Around 30-60% of the human population already has preexisting antibodies against most or all AAV serotypes.^7,8^ This triggers immunotoxicity and precludes clinical efficacy, making such patients ineligible for treatment. Finding ways to circumvent neutralizing antibodies represents a central challenge in the field of AAV gene therapy.

A handful of approaches for circumventing anti-AAV neutralizing antibodies have been attempted.^9^ Capsid engineering by directed evolution^10^ and machine learning approaches^11^ can yield antigenically distinct AAVs that evade antibodies to some level. But these methods are constrained by the locations of the capsid mutations, so some changes do not improve immune evasion and some changes reduce the infectivity of the virus.^9,12^ Multiple rounds of plasmapheresis coupled with pharmacological immunosuppression has shown some success, but this process is expensive, invasive, and risky.^13^ Another promising route arose after Maguire et al. demonstrated in 2012 that a fraction of AAVs coming out of producer cells are packaged into exosomes (exoAAVs) and that these complexes enjoyed partial protection from neutralizing antibodies.^14^ However, exoAAVs have encountered obstacles in the form of poor yields, purity issues, and difficulties in accurately determining titers for consistent therapeutic administration in animal models.^15^ Chemical modification of AAV capsids with polyethylene glycol has shown partial protection from neutralizing antibodies at the expense of substantially decreasing cellular transduction efficiency.^16^ Although some progress has been made, it thus remains clear that improved ways of overcoming neutralizing antibodies are desperately needed for AAV gene therapy to reach its full potential.

Vaults are ribonucleoprotein particles found in most eukaryotes and have been proposed to have been present in the last eukaryote common ancestor.^17–19^ The particle shell can be produced recombinantly using insect cells or yeast cells.^20,21^ These recombinant particles are structurally indistinguishable from native vaults except that they are hollow inside. Vaults are assembled on polyribosomes from 78 copies of major vault protein (MVP).^22^ These recombinant particles have previously been repurposed as drug delivery vehicles^23,24^ and as vaccines.^25–27^ Empty recombinant vaults are not immunogenic.^24,27^ Though vaults have a strong natural tropism for antigen presenting cells, activated T cells, and other phagocytic cells,^28^ they can be retargeted to transduce desired cell types via C-terminal modifications of MVP.^29^ Packaging of various protein and drug cargos into vaults has been achieved by fusing or conjugating the target molecule to INT, a peptide originally derived from a natural vault cargo protein known as PARP4,^23^ which binds the inside of the vault with high affinity.^23,30,31^ Of benefit for delivery applications, vaults disassemble during endosomal acidification, allowing cargo to escape the compartment.^31^ Interestingly, packaging of the adenoviral endosomolytic protein pVI into vaults has facilitated enhanced endosomal escape of ribotoxin cargo.^32^ Major difficulties have been encountered in packaging nucleic acids into vaults and strategies to overcome these issues have not been successful.^33^ Nonetheless, vaults have seen much use as a tool for delivery of small protein and drug cargos into cells, particularly in the context of intranasal vaccines.^25–27^

We herein develop a novel delivery system for AAVs which circumvents neutralizing antibodies from serum and conveys enhanced cellular transduction efficiency. Our technology leverages SpyTag-SpyCatcher technology^34,35^ to decorate AAV with INT peptide and thus facilitate its packaging inside recombinant vaults, resulting in vaultAAV (VAAV). To the best of our knowledge, no other group has packaged an entire virus into a vault. Remarkably, VAAV transduces cultured cells even in the presence of neutralizing serum. As such, our novel VAAV technology has high promise as a new way of overcoming the neutralizing antibody problem of AAV gene therapy.

## Results

As a way of site-specifically conjugating INT onto AAV, we leveraged the molecular glue peptides SpyTag-SpyCatcher. We genetically inserted SpyTag003 (SpT3) as flanked by flexible linkers into the VP2 protein of AAV9 **(Figure 2A)** and confirmed incorporation of VP2-SpT3 into the capsid by Western blot **(Figure 2B)**. Because each AAV capsid on average contains five copies of VP2 out of a total of sixty capsid proteins,^37^ we reasoned that this would minimally disrupt viral production and function. We next designed a SpyCatcher003-INT (SpC3-INT) fusion protein wherein a flexible linker connected the SpC3 and the INT peptide **(Figure 2A)**. We validated the SpC3-INT protein by SDS-PAGE **(Figure 2C)**. After this, we mixed AAV9-VP2-SpT3 with SpC3-INT to induce spontaneous formation of AAV9-VP2-SpT3-SpC3-INT, hereafter referred to as AAV9-INT.

**Figure 1.**
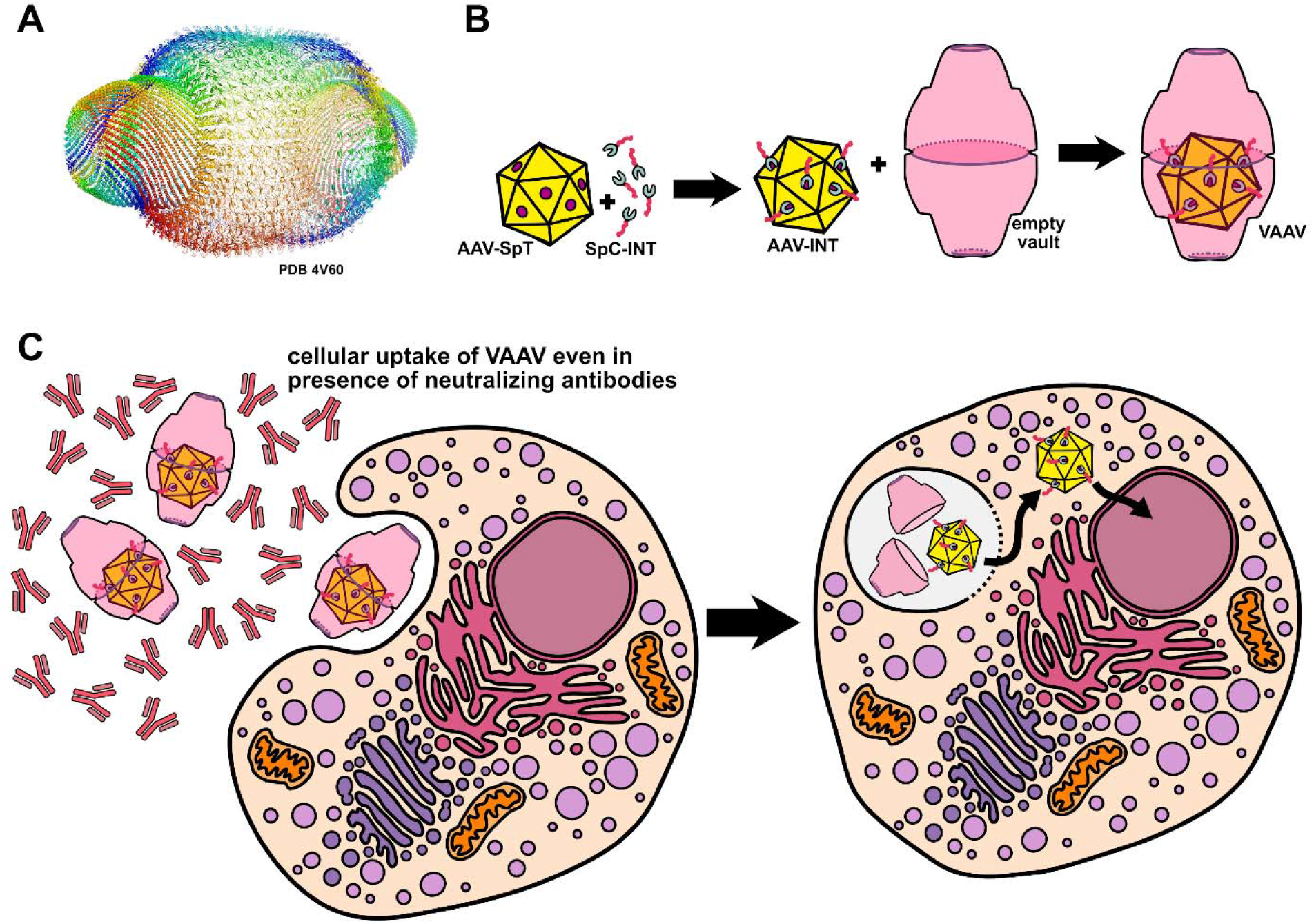
**(A)** Structure of the outer shell of a protein vault complex (PDB 4V60). The vault shell consists of 78 copies of MVP assembled into a hollow compartment. **(B)** We packaged AAV into recombinant vaults by (i) producing an AAV9 with SpyTag003 inserted into its VP2 capsid protein, (ii) creating a SpyCatcher003-INT fusion protein, (iii) conjugating the SpC3-INT with the AAV9-VP2-SpT3, and (iv) mixing the AAV-INT particles with empty vaults and incubating overnight to induce packaging. **(C)** Schematic of VAAV complexes transducing a cell despite the presence of anti-AAV neutralizing antibodies. The vault appears to behave as an immunologically invisible shield. After uptake, the vault disassembles within the acidifying endosome,^31^ the AAV exposes its VP1 phospholipase A2 domain to facilitate endosomal escape, and the AAV experiences transport to the nucleus.^36^

**Figure 2.**
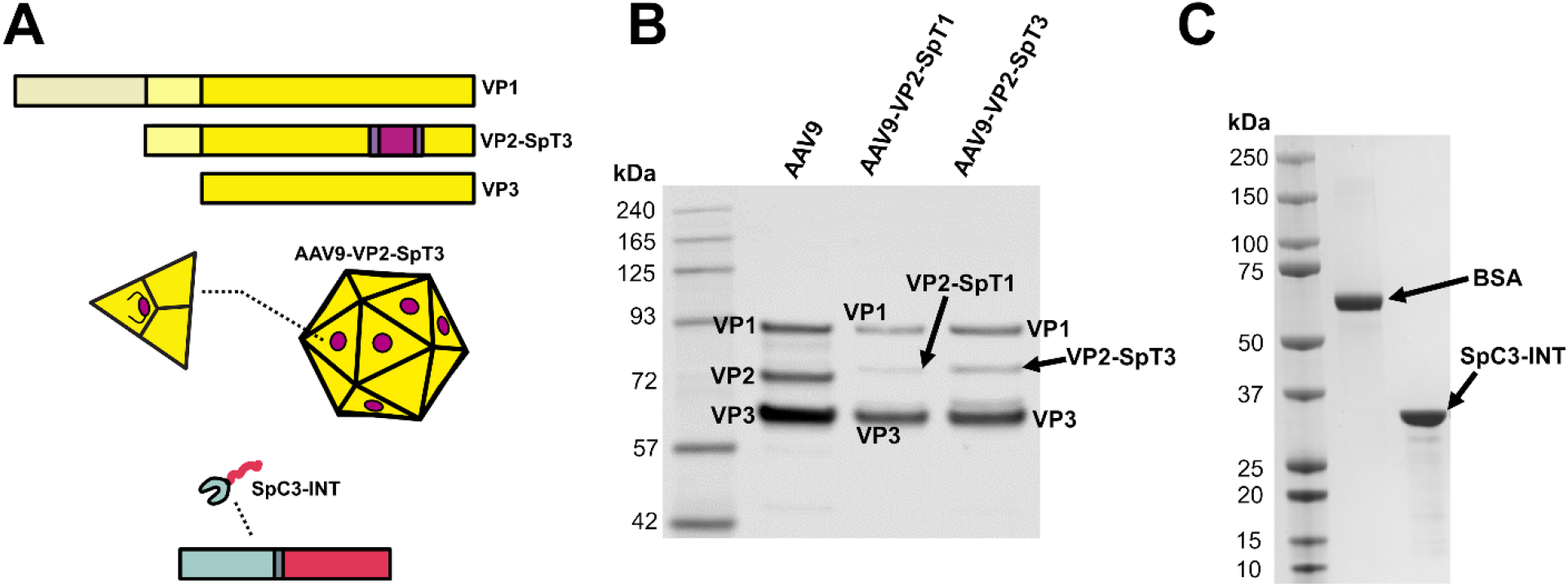
**(A)** AAV9-VP2-SpT3 design includes SpT3 flanked by flexible linkers and inserted into the VP2 capsid protein. To prevent the SpT3 from appearing in VP1 and VP3, we used two separate plasmids to encode (i) VP2-SpT3 and to encode (ii) VP1 and VP3 during AAV production. SpC3-INT was designed as a C-terminal fusion of INT peptide onto SpC3, bridged by a flexible linker. **(B)** Western blot of the AAV capsid proteins validated successful production of wild-type AAV9, AAV9-VP2-SpT1, and AAV9-VP2-SpT3. The VP2-SpT1 and VP2-SpT3 proteins show the expected ∼1 kDa shift on the gel relative to VP2. We included AAV9-VP2-SpT1 in the Western blot as an additional control, though we chose not to further utilize this virus in our study since SpT3 exhibits superior reaction kinetics.^35^ **(C)** SDS-PAGE of SpC3-INT confirmed its expected molecular weight after production.

We mixed AAV9-INT with purified vaults at a 1:5 particle count ratio and incubated them overnight at room temperature to facilitate packaging. Since vaults spontaneously package INT-bearing biomolecules,^23,24^ we anticipated they might be able to encapsulate AAV-INT despite its larger size and complexity compared to past vault cargos. We first imaged AAV and vaults alone to verify their morphological characteristics **(Figure 3A)**. After mixing AAV-INT with vaults, we were surprised to find that just overnight incubation at room temperature was sufficient to induce packaging of a sizable fraction of AAVs into vaults, though many free AAVs did remain unencapsulated **(Figure 3B-D, Figure S1)**. It should be noted that future optimization of this technique may improve the packaging frequency. Transmission electron microscopy (TEM) imaging validated formation of the VAAV complexes. Remarkably, a small number of VAAVs appeared to encapsulate two AAVs at once **(Figure 3B)**. By comparison, AAV9-VP2-SpT3 which had been reacted with SpC3 (without the INT peptide) did not undergo packaging into vaults **(Figure 3C)**. These novel technical results encouraged us to further develop VAAV as a gene therapy delivery system.

**Figure 3.**
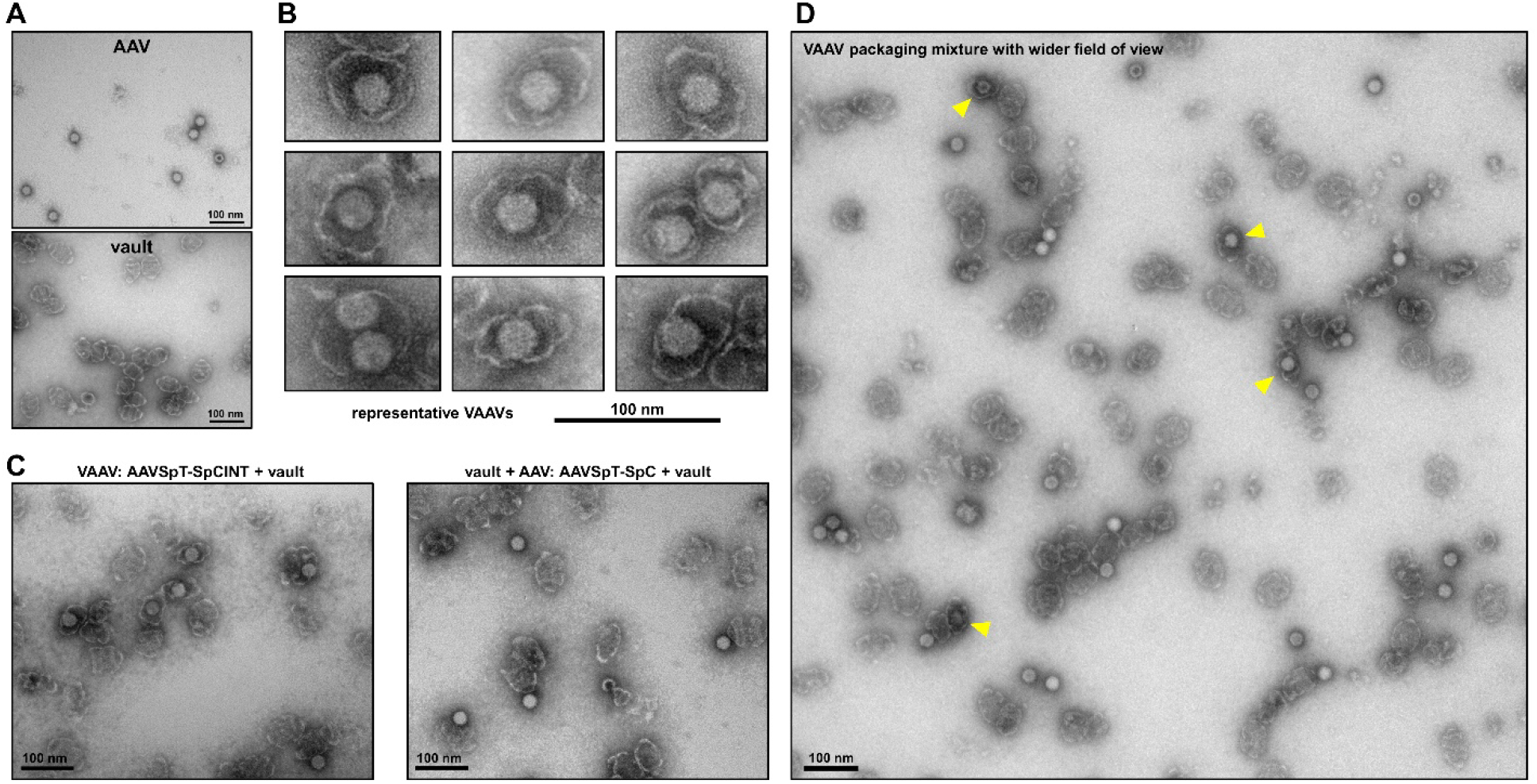
**(A)** TEM images of purified AAV and purified vault samples. **(B)** Representative examples of VAAV complexes. **(C)** TEM comparison of VAAV sample and control vault + AAV sample wherein the latter contains a mixture of vaults with AAV9-VP2-SpT3 conjugated to SpC3. In the VAAV sample, a substantial fraction of AAV-INT undergoes packaging by vaults. In the vault + AAV sample, the AAVs lack the INT peptide and thus do not experience packaging into vaults. **(D)** Wider field of view representative image shows that although VAAV packaging occurs with reasonably high frequency, many unpackaged AAVs remain present in the mixture. Here, packaged VAAVs are marked with yellow arrows.

To locate AAVs more precisely within vaults, cryogenic electron tomography (cryoET) was performed. Tomogram slices show the distribution of vaults and AAVs in three orthogonal directions **(Figure 4A)**. Three vaults with different orientations are recognizable in the slices. The hexagonal density (green arrows) with darker contrast comes from the icosahedral AAV, which resides exactly inside the central cavity of the vault from all three views. The 3D rendering of this VAAV particle **(Figure 4B)** shows that the AAV (green) is well encapsulated by the vault shell (purple). Vault structures shown in both tomogram slices and 3D rendering are not fully closed due to the missing wedge reconstruction issue, which made the whole density distort and elongate in the Z direction. These cryoET data give direct evidence that AAVs are indeed packaged inside of vaults.

**Figure 4.**
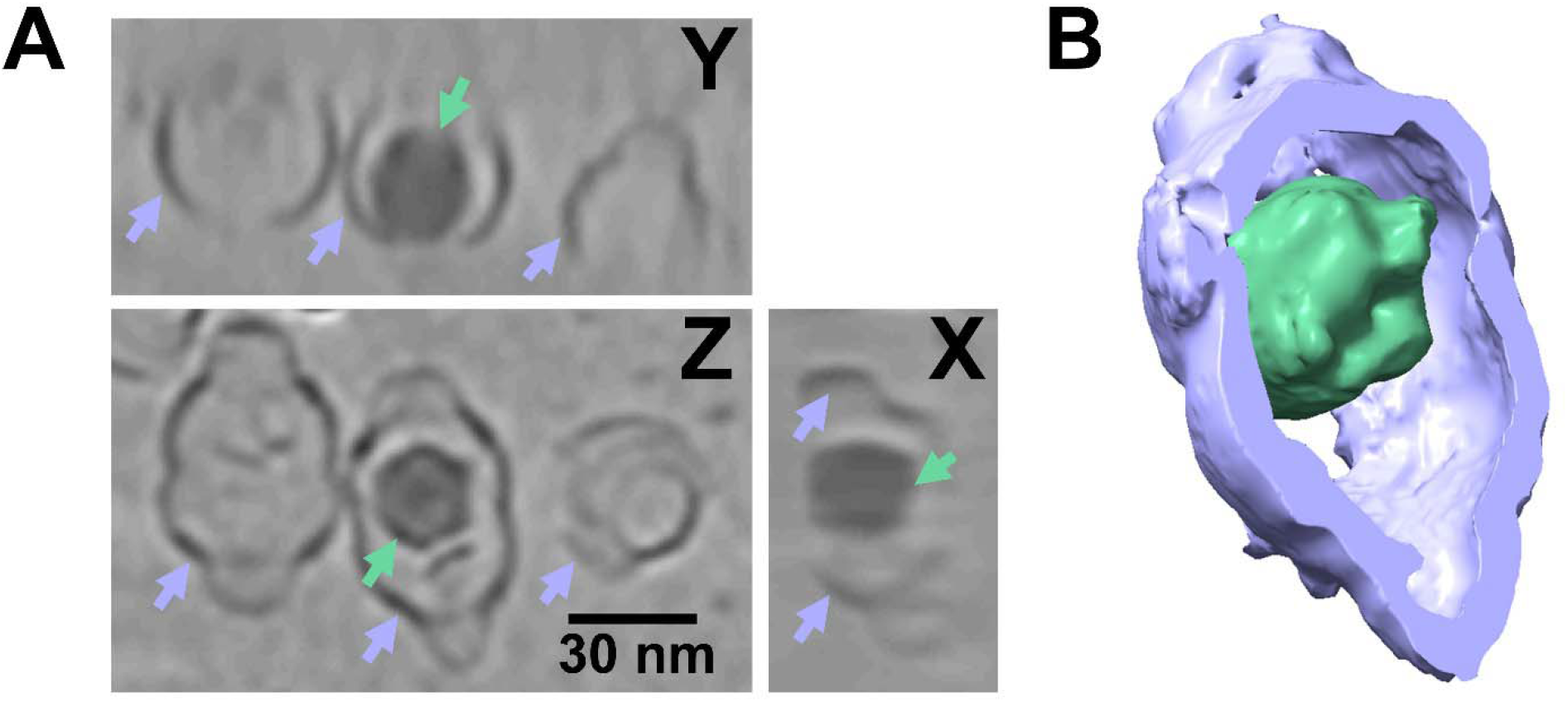
**(A)** Tomogram slices of VAAV and vault particles, showing the three views in orthogonal directions. Vaults and AAVs are marked with purple and green arrows respectively. **(B)** 3D rendering of a tomogram with a VAAV particle. Vault and AAV are colored in purple and green respectively. Note that the reason the vault structure is not fully closed is due to the missing wedge reconstruction issue rather than any actual damage to the vault’s wall.

As a proof-of-principle to test the immunological shielding capacity of VAAV, we utilized neutralizing serum from mice immunized against AAV9-VP2-SpT3. We first performed a neutralization assay to verify that the serum completely blocked AAV9-VP2-SpT3 transduction of cultured CHO-Lec2 cells **(Figure S2)**. Then we treated CHO-Lec2 cells with VAAV, vault + AAV, AAV9-INT alone, and PBS both in the presence and the absence of a fully neutralizing level of serum. Even with neutralizing serum, VAAV successfully transduced cells **(Figure 5A-B)**. Roughly speaking, the percentage of transduction paralleled the amount of packaged VAAVs we observed in the TEM images. By contrast, vault + AAV showed a very low level of transduction, AAV alone did not transduce cells, and no fluorescence was observed in the PBS group. In the absence of neutralizing serum, VAAV and vault + AAV very strongly transduced cells, AAV alone transduced cells to a moderate extent, and no fluorescence was observed in the PBS group **(Figure 5A, 5C)**. Enhanced vault + AAV transduction might arise from nonspecific associations between vault and AAV bringing AAV “along for the ride” when vault undergoes endocytic uptake into the cells. It is noteworthy that similar enhancement has been observed in context of vaults improving transfection of nonpackaged plasmids and has previously been termed the “bystander effect”.^24,32,38^ These data demonstrate that VAAV protects its AAV cargo from neutralizing serum as well as enhances cellular transduction, making it a promising new delivery vehicle for circumventing the neutralizing antibody problem of AAV gene therapy.

**Figure 5.**
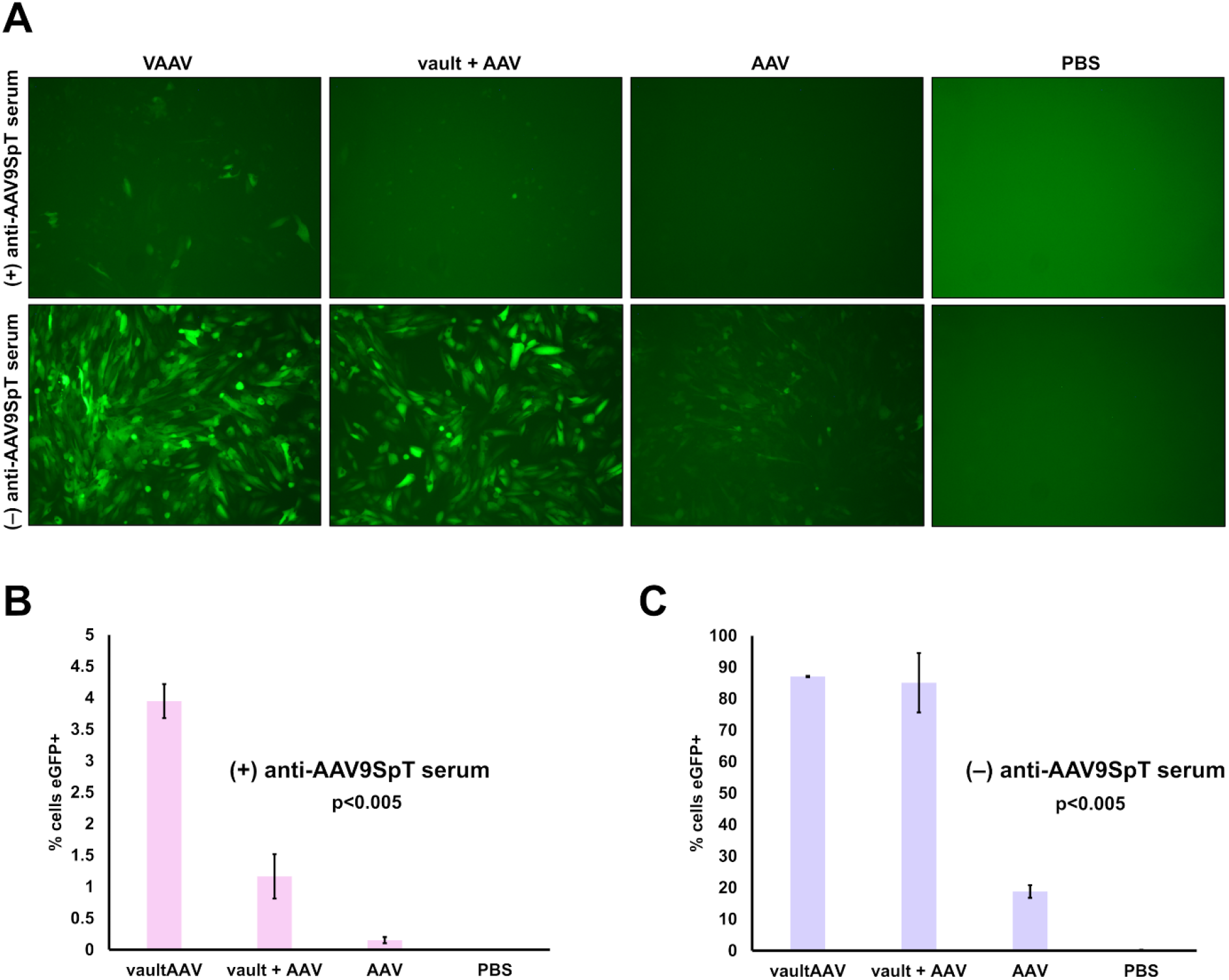
**(A)** Representative fluorescence microscopy images showing VAAV cellular transduction in the presence of neutralizing serum alongside controls. VAAV transduces cells in the presence of neutralizing serum, vault + AAV transduces cells to a low level in the presence of neutralizing serum, AAV alone does not transduce cells in the presence of neutralizing serum, and the PBS group shows no fluorescence. Both VAAV and vault + AAV very strongly transduce cells in the absence of neutralizing serum, AAV alone transduces cells to a moderate extent in the absence of neutralizing serum, and the PBS group shows no fluorescence. **(B)** Flow cytometric quantification of VAAV, vault + AAV, AAV, and PBS group transduction of cells in the presence of neutralizing serum. **(C)** Flow cytometric quantification of VAAV, vault + AAV, AAV, and PBS group transduction of cells in the absence of neutralizing serum.

## Discussion

AAV gene therapy has long been plagued by the neutralizing antibody problem and solutions have so far been limited.^9,12^ VAAV offers a promising new route for attacking this problem. VAAV protects AAV capsids via the immunologically invisible vault shell, facilitating successful delivery even when neutralizing antibodies are present. We anticipate that this platform technology will possess wide applicability since even though vault has a natural tropism for phagocytic cells, one can retarget the complex by attaching peptides or antibodies.^29,39^ Our proof-of-principle data showcase the potential of VAAV for circumventing the neutralizing antibody problem.

In addition, we show that mixtures of vaults and AAVs exhibit greatly enhanced cellular transduction efficiency regardless of whether AAVs are packaged into vaults. This might arise from vaults nonspecifically associating with AAVs and thus taking them along during endocytosis or the previously mentioned “bystander effect”.^32,38^ Enhanced VAAV transduction might help alleviate the need for utilizing potentially toxic high doses of AAV.^5^ Lower doses of AAV also may ease manufacturing burden and thus decrease the presently high costs of treatments.^1^ The strength of the effect in CHO-Lec2, a cell type known to endocytose vaults with a relatively modest degree of efficiency,^39^ supports the idea that the effect may not depend on cell type-specific mechanisms. Cell types which endocytose vaults more efficiently (e.g. phagocytes) may thus show the same or an even stronger transduction enhancing effect. It should be noted that this will likely still necessitate creating an AAV-SpT3 serotype which experiences efficient trafficking to the nucleus of the target cell type after endocytic uptake. VAAV’s enhanced transduction is an exciting property that will benefit from further exploration.

Despite vault’s tropism for phagocytic cells,^28^ past investigations have shown that it can be retargeted to transduce a variety of useful tissues.^25,29,39^ As an example, Kickhoefer et al. linked an epidermal growth factor (EGF) peptide to the C-terminus of MVP within vaults, successfully targeting uptake into epithelial cancer cells.^29^ They also demonstrated that fusion of an Fc-binding protein to MVP along with addition of an anti-EGFR antibody enabled targeting of epithelial cancer cells. These modified vaults experienced minimal uptake into non-target cell types. As such, VAAV should be amenable to the same sorts of powerful targeting techniques. VAAV thus might gain footing as a highly versatile platform technology with capacity for both immunological shielding and tissue-specific targeting, making it useful for a wide variety of biomedical applications.

VAAV offers a potential new solution to the neutralizing antibody problem of AAV gene therapy as well as improves cellular transduction efficiency **(Figure 1A-C)**. This study provides a concrete proof-of-principle to demonstrate the promise of our technology. We employ SpyTag-SpyCatcher molecular glue peptides as a method of site-specific conjugation for the INT peptide **(Figure 2A-C)**, which binds to the interior of the vault and facilitates packaging. We image VAAV complexes via TEM and show that a large fraction of AAVs package inside of vaults **(Figure 3A-C)**. We furthermore verify that AAVs are packaged within the vault interior by using cryoET **(Figure 4A-B)**. AAV9-INT represents the largest cargo and the first virus ever packaged inside of a vault. Finally, we show that VAAV transduces cells in the presence of neutralizing serum and that VAAV in general exhibits a transduction efficiency enhancing effect **(Figure 5A-C)**. We anticipate VAAV could expand the scope of AAV medicines to treat patient populations who have preexisting neutralizing antibodies and might eventually enable redosable AAV therapeutics.

## Methods

### Acquisition of anti-AAV9-VP2-SpT3 serum and of SpC3-INT

Intramuscular injection of 5×10^11^ vg of AAV9-VP2-SpT3 was performed on C57BL/6 mice to induce production of anti-AAV-VP2-SpT3 serum. Serum was recovered and used in subsequent experiments. Neutralization was validated by incubating varying dilutions of serum with 10^6^ vg of AAV9-VP2-SpT3 for 1 hour at room temperature before transferring each of the samples onto 10^4^ CHO-Lec2 cells in a 96-well plate (10^6^ multiplicity of infection or MOI). At 4 days post-infection, fluorescence microscopy was utilized to observe what concentration of serum was necessary for full neutralization. As a custom service, GenScript produced SpC3-INT and validated it using SDS-PAGE.

### Production of AAV9-VP2-SpT3

AAV Rep-Cap plasmids containing the *Rep* gene from AAV2 and the *Cap* gene for AAV9 were produced by gene synthesis and molecular cloning. Plasmid pVP13 (plasmid identification number 1886) contains a mutation in the ACG start codon of VP2 to GCG so as to only enable expression of VP1 and VP3. Plasmid pVP2-SpT3 (plasmid identification number 3007) contains ATG to CTG mutations at the start codons of both VP1 and VP3 so as to express only the SpyTag003-modified VP2. The following amino acids were inserted after glycine 453 in the *Cap* sequence: flexible linker [AGGGSGGS], SpyTag003 [RGVPHIVMVDAYKRYK], flexible linker [GGSGGSA]. Transgene plasmid pAAV-CB-EGFP (plasmid identification number 1161) (PMID: 34141821) expresses enhanced greed fluorescent protein driven by the Chicken Beta Actin promoter with a CMV enhancer element. A plasmid expressing the Adenoviral helper genes required for AAV packaging (pAdDeltaF6) was obtained from the University of Pennsylvania Vector Core. AAV9-VP2-SpT3 were produced through quadruple transfection of 293T cells (ATCC, CRL-3216) using polyethylenimine (PEI) (PMID: 23791963). Transgene, helper, and Rep-Cap plasmids were supplied in an equimolar (1:1:1:1) ratio. At 72 hours post-transfection, cell pellets were harvested and recombinant AAV were purified by iodixanol gradient ultracentrifugation. Fractions containing AAV genomes were identified by qPCR, pooled, and dialyzed against PBS using a 100 kDa Spectra-Por Float-A-Lyzer G2 dialysis device (Spectrum Labs, G235059). Purified AAV were concentrated using a Sartorius Vivaspin Turbo 4 Ultrafiltration Unit (VS04T42) and stored at –80C. AAV titers were calculated by qPCR relative relative to a standard curve of transgene plasmid, after DNase digestion to remove unencapsidated DNA [PMID: 23912992]. The following primers were used to detect the EGFP transgene for titer: for 5’-GCATCGACTTCAAGGAGGAC-3’, rev 5’-TGCACGCTGCCGTCCTCGATG-3’.

### Production of vault

Codon-optimized DNA encoding human MVP was inserted into vector pJGG (BioGrammatics, San Diego, CA) and transformed into yeast (*Pichia pastoris*) cells. MVP-expressing *P. pastoris* was cultured in YPD medium at 30°C in shake flasks at 200 rpm for 24 hours. Harvested cells were resuspended in Buffer (50 mM Tris pH 7.4, 75 mM NaCl, 0.5 mM MgCl_2_) supplemented with 1 mM DTT and protease inhibitor cocktail (Sigma-Aldrich P8215) and lysed by agitating with 0.5 mm glass beads. Cellular debris and beads were removed by centrifugation at 3000 g for 5 minutes followed by centrifugation at 20,000 g for 20 minutes. Supernatant was further centrifuged at 37,000 rpm for 1 hour, and the vault-containing pellet was resuspended in Buffer containing 1% Triton X-100. Vaults were precipitated by addition of 0.1 g ammonium acetate per mL and agitated overnight at 4°C. Precipitate was collected by centrifugation at 20,000 g for 20 minutes, resuspended in Buffer, and remaining debris was removed by centrifugation at 3000 g for 10 minutes. Supernatant was applied to a TMAE Fractogel HiCap column (EMD) and eluted via NaCl gradient. Vault-containing fractions were pelleted at 37,000 rpm for 1 hour and resuspended to a concentration of 1 mg/mL in phosphate-buffered saline containing 10 mg/mL trehalose. Aliquots of resuspended vaults were frozen in liquid nitrogen, lyophilized, and stored at –20°C.

### Packaging of AAV into vault

For the VAAV group, AAV9-VP2-SpT3 was mixed with SpC3-INT at a 1:10 copy number ratio so that the average of 5 SpT3 sites per capsid would be saturated while not having such an excess as to crowd the interiors of the vaults in the next step. For the vault + AAV control, AAV9-VP2-SpT3 was mixed with the same amount of SpC3. For the AAV control, another aliquot of AAV9-VP2-SpT3 was mixed with the same amount of SpC3-INT. The mixtures were incubated for 1 hour at 25°C. For the VAAV group and the vault + AAV control, vault was then added to at 5-fold copy number excess relative to AAV particles. Number of vaults per volume was calculated based on molecular weight of a single vault and total vault mass resuspended in PBS with 10% glycerol. For the AAV control, an equivalent volume of PBS with 10% glycerol was added. All three mixtures were incubated at 25°C overnight. To validate packaging, a small amount of each group was removed, diluted 1:30 in PBS with 10% glycerol, and imaged via negative stain (1% uranyl acetate) TEM. The exact number of AAV particles removed from each sample for this validation was calculated to ensure accurate MOI calculations for the subsequent experiment.

### CryoET and tomogram reconstruction

To begin the cryoET process, 10 nm fiducial gold beads were added to the VAAV-containing sample. An aliquot of 3.5 μL of the mixture was briefly applied onto Quantifoil holey carbon grids (300 mesh, 1.2/1.3) with a 2 nm continuous carbon layer that was glow-discharged for 60 s at –25 mA with PELCO easiGlow. After a 2-minute waiting time, the extra sample on the grids was blotted away by filter paper and the resulting cryoEM grid was placed in an FEI/Thermo-Fisher Mark IV Vitrobot cryo-sample plunger at blot force 2 and blot time 3 s. The grids were next plunged into precooled liquid ethane/propane mixture for vitrification. Plunge-freezing conditions and the concentrations of vaults and AAVs were optimized by screening with a FEI TF20 TEM equipped with a Gatan K3 camera.

Tilt series were collected using *SerialEM* in a FEI Titan Krios transmission electron microscope equipped with an energy filter and a Gatan K3 camera at super-resolution mode.^40^ The collecting magnification was 64 kx and the tilt range was –60° to 60° with a 3° tilt-series increment and dose symmetry.^41^ Total electron exposure for one tilt series was 120 e^-^/Å^2^ with frame fraction. Frames in the tilt series were drift-corrected using *MotionCor2*^42^ and the corrected micrographs were aligned and reconstructed into tomograms with *AreTomo*.^43^ *IsoNet* was applied to enhance the contrast and partially alleviate the missing wedge issue.^44^ Visualization of tomogram slices and three-dimensional rendering of tomograms was performed using *IMOD*^45^ and *UCSF ChimeraX*.^46^

### Infection assay with and without neutralizing serum

Groups including VAAV, vault + AAV, AAV, and PBS were prepared and validated by TEM (as described in the previous section) prior to the infection experiment. CHO-Lec2 cells were seeded at 10^4^ cells per well in a 96-well plate. For the groups with neutralizing serum, a 1:100 dilution of serum was mixed with VAAV, vault + AAV, AAV, and PBS samples 1 hour before transferring the mixtures to appropriate wells in triplicate at an MOI of 10^6^ vg per cell. (Samples were incubated at room temperature during this 1-hour period). The final dilution of serum in the wells was 1:200. For the groups without neutralizing serum, mixtures of VAAV, vault + AAV, AAV only, and PBS were prepared and added to the appropriate wells in triplicate at an MOI of 10^6^ vg per cell. At 4 days post-infection, the cells were imaged by fluorescence microscopy. Flow cytometry was then employed to quantify the percentage of eGFP-positive cells in each well. A one-way ANOVA (analysis of variance) test was used to determine statistical significance.

## Acknowledgements

We thank Dr. Valerie Kickhoefer for her helpful advice on vault production and handling.

