## Supplementary Information for "Encapsulation of AAVs into protein vault nanoparticles as a novel solution to gene therapy’s neutralizing antibody problem"

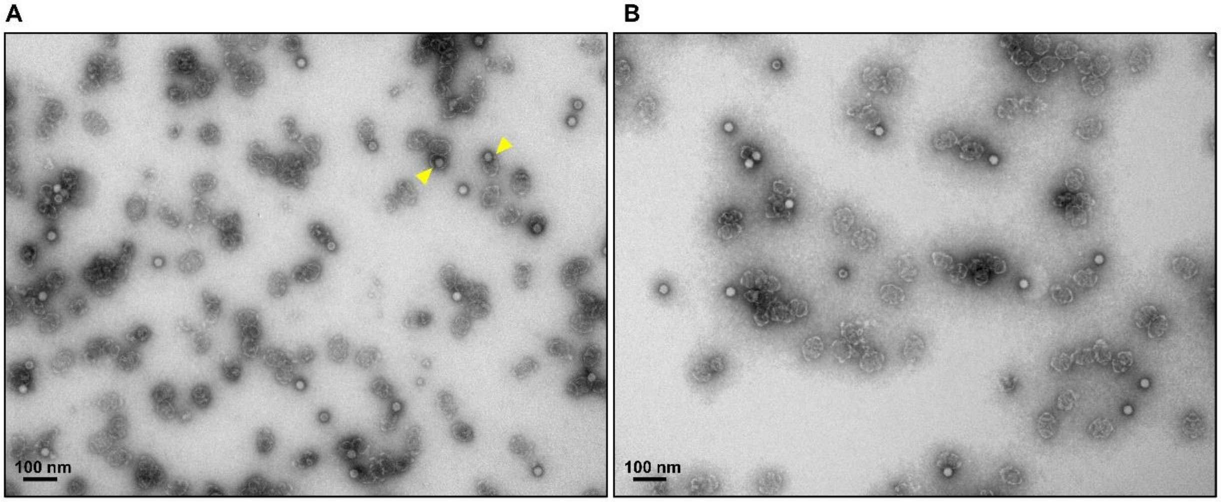


**Figure S1 (A)** Another representative larger field of view showing VAAV packaging mixture where a substantial fraction of AAV-INT has been packaged into vaults, yet a number of unpackaged AAVs still persist in the mixture. Here, packaged VAAVs are marked with yellow arrows. Ambiguous cases were not counted. **(B)** Representative larger field of view showing vault + AAV mixture where the AAVs did not package into vaults.

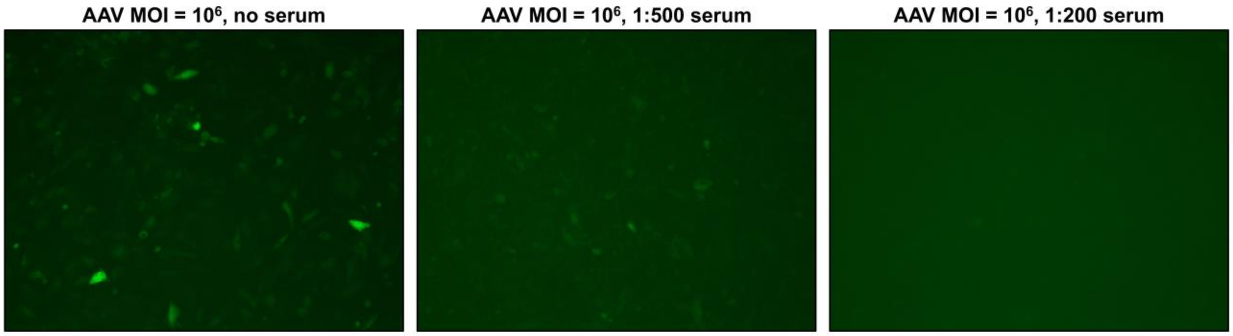


**Figure S2** Neutralization assay showing that anti-AAV9-VP2-SpT3 serum prevents AAV9-VP2-SpT3 ($10^6$ MOI) from transducing CHO-Lec2 cells at a 1:200 final serum dilution.
